# TACSTD2 upregulation is an early reaction to lung infection

**DOI:** 10.1101/2021.06.29.450320

**Authors:** Sára Lenárt, Peter Lenárt, Lucia Knopfová, Hana Kotasová, Vendula Pelková, Veronika Sedláková, Vladimír Čan, Jan Šmarda, Karel Souček, Aleš Hampl, Petr Beneš

## Abstract

*TACSTD2* encodes a transmembrane glycoprotein Trop2 commonly overexpressed in carcinomas. While the Trop2 protein was discovered already in 1981 and first antibody-drug conjugate targeting Trop2 were recently approved cancer therapy, the physiological role of Trop2 is still not fully understood. In this article, we show that TACSTD2/Trop2 expression is evolutionarily conserved in lungs of various vertebrates. By analysis of publicly available transcriptomic data we demonstrate that *TACSTD2* level consistently increases in lungs infected with miscellaneous pathogens. Single cell and subpopulation based transcriptomic data revealed that the major source of *TACSTD2* transcript are lung epithelial cells and their progenitors and that *TACSTD2* is induced directly in lung epithelial cells following infection. This increase may represent a mechanism to maintain/restore epithelial barrier function and contribute to regeneration process in infected/damaged lungs.

## Introduction

Trop2 is a transmembrane glycoprotein with yet unresolved physiological function, that is overexpressed in most carcinomas where it has been associated with cancer cell plasticity, tumor growth, metastasis and prognosis (1,2). It is encoded by the intronless *TACSTD2* (tumor-associated calcium signal transducer 2) gene belonging to *TACSTD* gene family (3). Genes of the *TACSTD* gene family are highly conserved across species; for instance, mouseTrop2 is 79.2% identical and 87.4% similar to human Trop2 (4,5). Trop2 was originally found on the surface of trophoblast cells (6) and has been subsequently identified on healthy epithelial cells of various other organs (7,8). Trop2 is also expressed during normal embryonal and fetal development in lungs (9,10), intestines (11), stomach (12), urinary bladder (13), kidneys (14), and cerebellum (15), however, its function in healthy adult tissues remains unknown.

In humans, congenital mutations of *TACSTD2* cause a gelatinous drop-like corneal disease (GDLD), a rare autosomal recessive disease characterized by the development of bilateral corneal amyloidosis and eventually blindness (16). Loss of the Trop2 function leads to impaired subcellular localizations of tight junction-related proteins and loss of barrier function of corneal epithelial cells resulting in passage of Lactoferrin to subepithelial region where it forms amyloid deposits (17). Trop2 is also considered to be a stem/progenitor cell marker (11,13,18–21) and several studies indicate that it might be associated with tissue remodeling and regeneration processes (12,22,23). Surprisingly, the *Tacstd2* null mice are fully viable, fertile, and without overt developmental abnormalities (24).

Lungs are vital organs inherently vulnerable to infection and injury due to constant exposure to pathogens, chemicals, and other air pollutants. The proper functions of epithelial barrier, immune system, and regenerative capacity of the lungs are thus crucial for restoring homeostasis following pathogen exposure or acute injury (25). The importance of lung homeostasis maintenance is further highlighted by the fact that even before the rise of SARS-CoV-2 pandemic, respiratory diseases belonged to leading causes of death worldwide (26). In this study, we use available expression datasets to test the hypothesis that the upregulation of *TACSTD2* in the lungs is a physiological reaction to infection or injury, which both trigger an acute immune response (27–29).

## Methods

### Lung expression data analysis

In order to analyze *TACSTD2*/Trop2 expression in healthy lungs, several databases were searched. First, “TACSTD2” was searched in every species included in the Expression atlas database (30) (https://www.ebi.ac.uk/gxa/home, accessed on January 5, 2021). The results were filtered for baseline lung expression in each organism. From each study, information about expression level and the number of biological replicates was retrieved. The expression value was set to 0.5.

“TACSTD2” was also searched in The Human Protein Atlas (31) (https://www.proteinatlas.org/, accessed on January 5, 2021) where the Tissue atlas was selected, and Lung was chosen to obtain RNA and protein expression data.

“TACSTD2” was also searched in GTEx Portal (32) (https://www.gtexportal.org/home/, accessed on January 5, 2021, dbGaP accession number: phs000424.v8.p2). Even though, data we used from GTEx are also available in Expression Atlas and The Human Protein Atlas, the GTEx is the primary source and data in other databases are not always up to date. Therefore, when data were available from multiple sources, we used data from GTEx for analyses.

Furthermore, “TACSTD2” was searched in Bgee database (33) (https://bgee.org/, accessed on January 5, 2021). Because Bgee database uses ArrayExpress, GEO, and GTEx – dbGAP as sources of raw data, and the same sources are also used by the Expression Atlas, *TACSTD2* expression data from Bgee were used only for animals which *TACSTD2* expression profile was not listed in Expression Atlas database, i.e., pig, chimpanzee, macaque, rabbit, and opossum (see Suplementary Data S1).

### Differential expression data analysis

To test the hypothesis that the overexpression of *TACSTD2* in the lungs is a physiological reaction to infection or injury, we first analyzed differential *TACSTD2* expression data for every species included in the Expression atlas database (30) (https://www.ebi.ac.uk/gxa/home, accessed on January 5, 2021). Results were filtered by choosing Infect or Injury in Experimental variables. Subsequently, only studies that met the following criteria were included:

1. The dataset must contain data from experiments with whole organisms, not cell lines.
2. The dataset provides information about lung transcriptome.
3. The dataset allows identification of differentially expressed genes following infection or injury.

Additionally, the E-GEOD-33266 dataset, which was not found by the above-described approach as it did not contain the keyword “Infect” in its annotation, was identified by searching Expression Atlas COVID-19 Data Portal.

In each particular study, Log_2_-fold change value was set to 0.0 so we could see even the small change of *TACSTD2* expression. The adjusted p-value was set to 1, so we could see non-significant results as well. P-values were adjusted for multiple testing using the Benjamini and Hochberg false discovery rate (FDR) correction (34).

In order to pinpoint cell population(s) responsible for the increase of *TACSTD2*/Trop2 after infection in the lungs, we searched for datasets in Gene Expression Omnibus (GEO) (35) containing sorted cell populations from infected lungs or infected cell cultures *in vitro*. Expression values for *TACSTD2/Tacstd2* from indicated datasets were retrieved, plotted and analyzed using GraphPad Prism v6.07 and shown in heat map using FGCZ Heatmap tool (http://fgcz-shiny.uzh.ch).

### Immunohistochemistry

The sample of human lungs was obtained from therapeutical surgery based on the written informed consent by the patient and approval of Ethics Committee of the University Hospital Brno (28-170621/EK). The specimens of pig, rat, mouse and human lungs were fixed with formalin, washed with PBS, dehydrated through series of alcohols (70%, 80%, and 96%) and embedded in paraffin blocks. Sections (4 μm thick) were dewaxed in xylene, hydrated through a graded series of alcohols (96%, 80%, and 70%), and rinsed in deionized water. After antigen retrieval in citrate buffer (pH 9.0, 12 min) at 98 °C for 30 min., the slides were rinsed in tap and deionized water and washed with 3% H_2_O_2_ in PBS at room temperature (RT) for 10 min. To block endogenous peroxidase activity, the sections were treated with 10% fetal bovine serum for 30 min. The sections were incubated with the primary antibody against TROP2 (Abcam, ab227689, 1:100) for 1 h at RT. The slides were then washed three times in PBS and subsequently incubated with the secondary antibody (En Vision FLEX/HRP) for 20 min. After the last washing step, the slides were incubated in substrate solution (DAB), counterstained in hematoxylin, dehydrated with alcohols and xylene, and mounted.

## Results

### *TACSTD2/Trop2* expression in lungs

To test the hypothesis that upregulation of *TACSTD2*/Trop2 is a physiological reaction to lung tissue damage by infection or injury, we first verified that *TACSTD2* is indeed expressed in healthy lungs. Analysis of available datasets shows overwhelming evidence that the *TACTSD2* gene is expressed in lungs of all studied species (**Table 1**). This suggests that *TACSTD2* has an evolutionarily conserved role in the function of lungs. A more detailed table is available in Supplementary data S1.

**Table 1.**
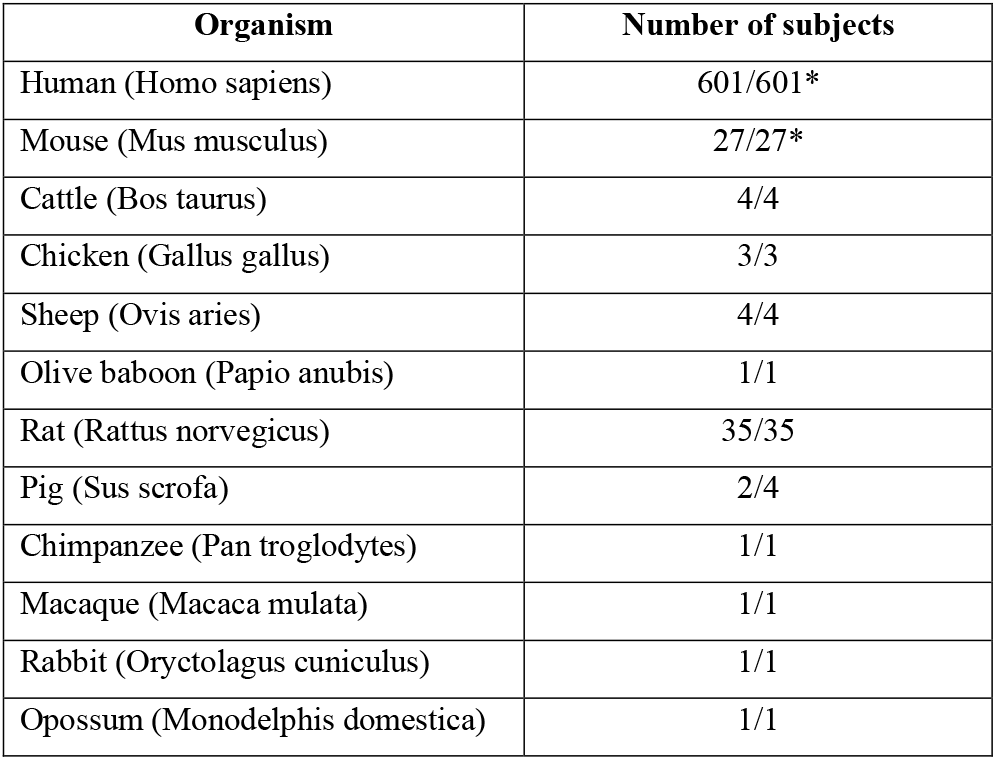
List of studied organisms with TACSTD2 expression in lungs. The number of subjects reflect the sum of biological replicates from transcriptomic datasets. * Trop2 expression was also found in human lungs in two of three proteomics datasets. The level of mouse Trop2 was below the cutoff in the one available proteomic dataset (see Supplementary data S1).

| Organism | Number of subjects |
| --- | --- |
| Human (*Homo sapiens*) | 601/601* |
| Mouse (*Mus musculus*) | 27/27* |
| Cattle (*Bos taurus*) | 4/4 |
| Chicken (*Gallus gallus*) | 3/3 |
| Sheep (*Ovis aries*) | 4/4 |
| Olive baboon (*Papio anubis*) | 1/1 |
| Rat (*Rattus norvegicus*) | 35/35 |
| Pig (*Sus scrofa*) | 2/4 |
| Chimpanzee (*Pan troglodytes*) | 1/1 |
| Macaque (*Macaca mulata*) | 1/1 |
| Rabbit (*Oryctolagus cuniculus*) | 1/1 |
| Opossum (*Monodelphis domestica*) | 1/1 |

To find which cell types produce *TACSTD2* in the lungs, we searched the Human Protein Atlas. The highest expression has been detected in alveolar cells type I and II, club cells and ciliated cells but smaller amounts of *TACSTD2* were also expressed in lung’s immune cells such as macrophages, T-cells, and granulocytes (**Figure 1A**). Recently, single cell transcriptomic analysis revealed that out of 58 molecular cell types identified in human lungs, *TACSTD2* is enriched in basal, differentiating basal, proliferating basal, proximal basal, goblet, alveolar epithelial type 1, platelets, and myeloid dendritic cells (**Figure 1B**) (36).

**Figure 1.**
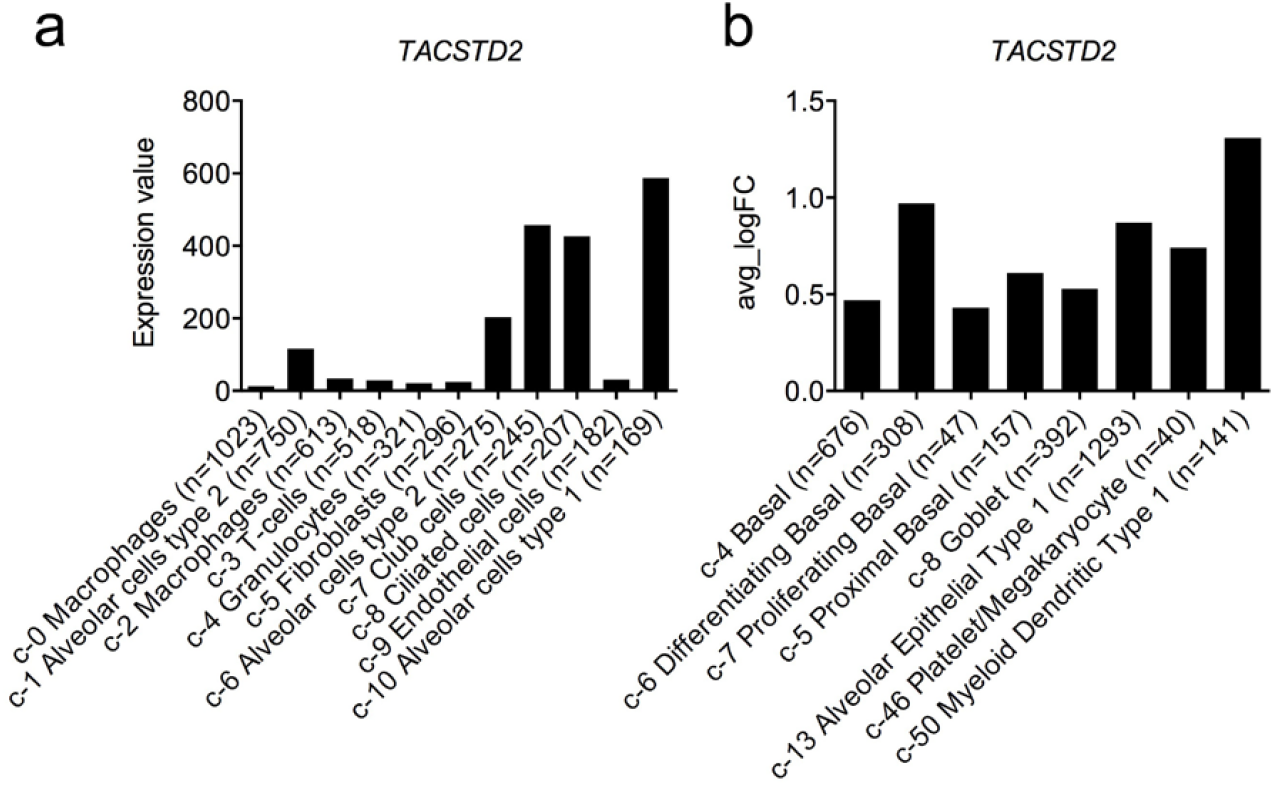
TACSTD2 expression in cell clusters of human lungs identified by single cell RNAsequencing. **A)** RNA expression (pTPM) in the cell type clusters identified in lung visualized by a bar chart, retrieved from Human Protein Atlas. Single cell transcriptomic dataset of Vieira Braga FA et al. (2019) (GSE130148) (37) was used. **B)** Cell clusters with significantly (p < 0.05) enriched TACSTD2 expression as identified in human lungs by Travaglini et al. 2020 (36). Chart shows the natural log of the average fold change between the indicated cell type and other cell types in lungs. c = cluster number, with main cell type annotated, n = number of included cells.

Interestingly, one mouse (E-MTAB-3579) and one rat (E-GEOD-53960) (38) dataset evaluated transcripts at different stages of embryonal development and at different stages of postnatal life. In these datasets, *Tacstd2* expression increased with age (**Figure 2A and 2C**). This has been recently confirmed by Angelidis *et al*., who detected significantly higher *Tacstd2* mRNA in the bulk lung RNA of 24-months-old mice than in the lungs of 3-months-old mice (**Figure 2B**). Single cell transcriptomic approach, however, did not reveal the source of this increase (39).

**Figure 2.**
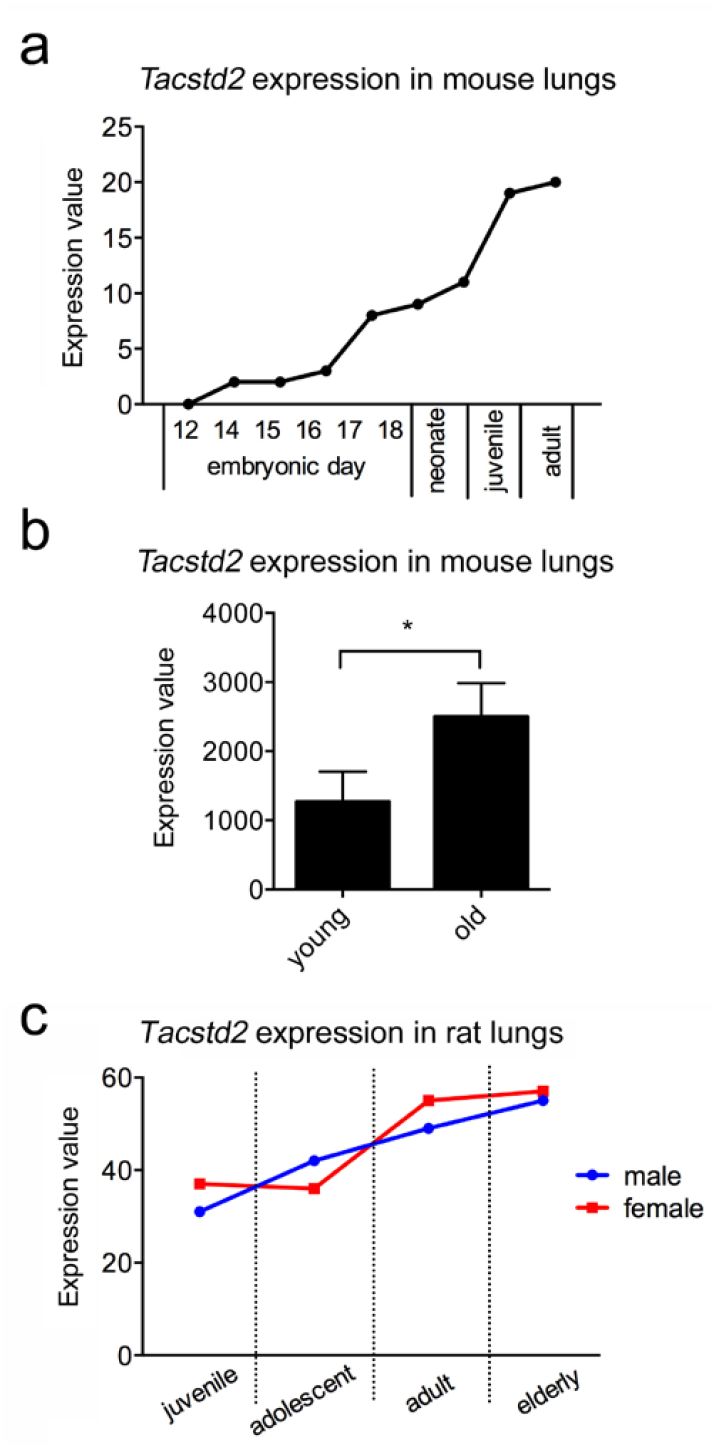
Tacstd2 gene expression in lungs (bulk data) of **A)** six mice during embryonal development, five neonate, two juvenile mice and one adult mouse (E-MTAB-3579). **B)** three replicates of young (3 months) and old mice (24 months) (39). **C)** juvenile (2 weeks), adolescent (6 weeks), adult (21 weeks) and elderly (104 weeks) female and male rats (E-GEOD-53960) (38). Four biological replicates were used for each sex and developmental stage.

To confirm that Trop2 protein is expressed in lungs of various organisms we performed IHC analysis in paraffin sections of human, mouse and pig lung tissues. In human lungs, Trop2 staining was observed in membranes of airway and alveolar epithelial cells while only basolateral parts of airway epithelium was Trop2 positive in mouse and pig lungs (**Figure 3**, Supplementary Figure S1). These data confirms that Trop2 is produced by lung epithelial cells but also point to differences in its expression pattern in lungs of selected vertebrates.

**Figure 3.**
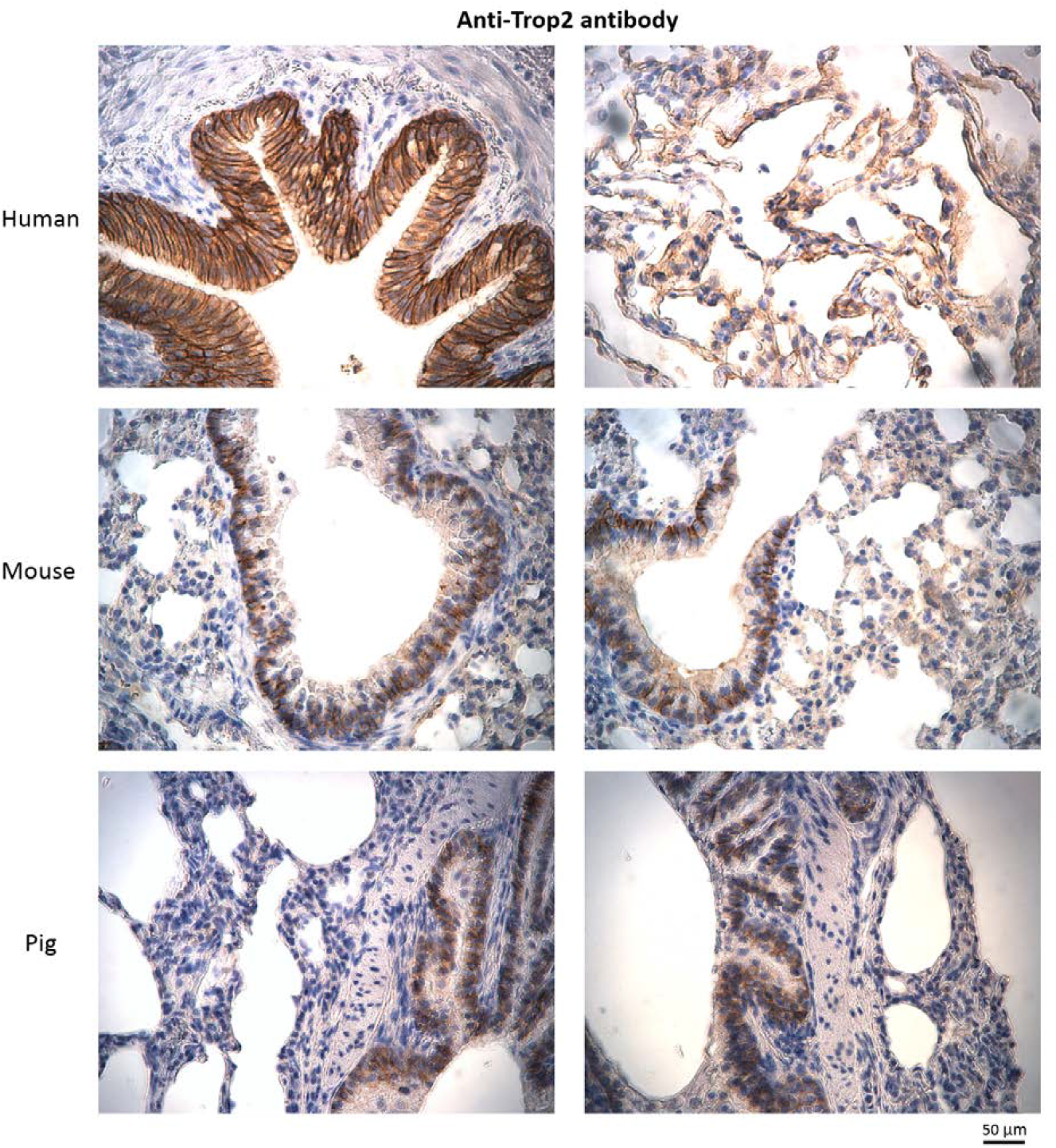
Immunohistochemical detection of Trop2 in paraffin sections of human, mouse and pig lung tissue. Human lungs – positive staining in epithelium of airway and alveoli. Mouse/Pig lungs – positive staining only in basolateral parts of airway epithelium.

### Upregulation of *TACSTD2* in infected lungs

Next, to analyze *TACSTD2* expression in response to lung damage, we searched the Expression Atlas database for differential expression in lungs after infection or injury. Eleven differential expression datasets analyzed the levels of *Tacstd2* early after infection (1-7 days), and all of them showed a significant upregulation of *Tacstd2* in infected mouse lungs (**Table 2**). This increase was observed in both males and females, 9 different mouse strains, at various ages (8-20 weeks), and with various infectious agents. Datasets containing information about the dynamics of this process showed that in the case of SARS coronavirus MA15, the levels of *Tacstd2* peaked two days post-infection and then started to decrease, indicating that the upregulation of *Tacstd2* is an early reaction to infection (**Table 2, Figure 4**). After infection with the influenza A virus, *Tacstd2* levels peaked on the fourth day, but the increase rate was reduced after the second day confirming that *Tacstd2* elevation is an early response (**Table 2**). Particularly intriguing is the E-GEOD-33266 dataset, where various doses of SARS coronavirus were used for infecting mice. Analysis of this dataset not only confirmed that *Tacstd2* levels were highest two days after SARS coronavirus infection regardless of the infection dose but also showed that a higher dose of the virus induced higher upregulation of *TACSTD2* **(Table 2)**.

**Figure 4.**
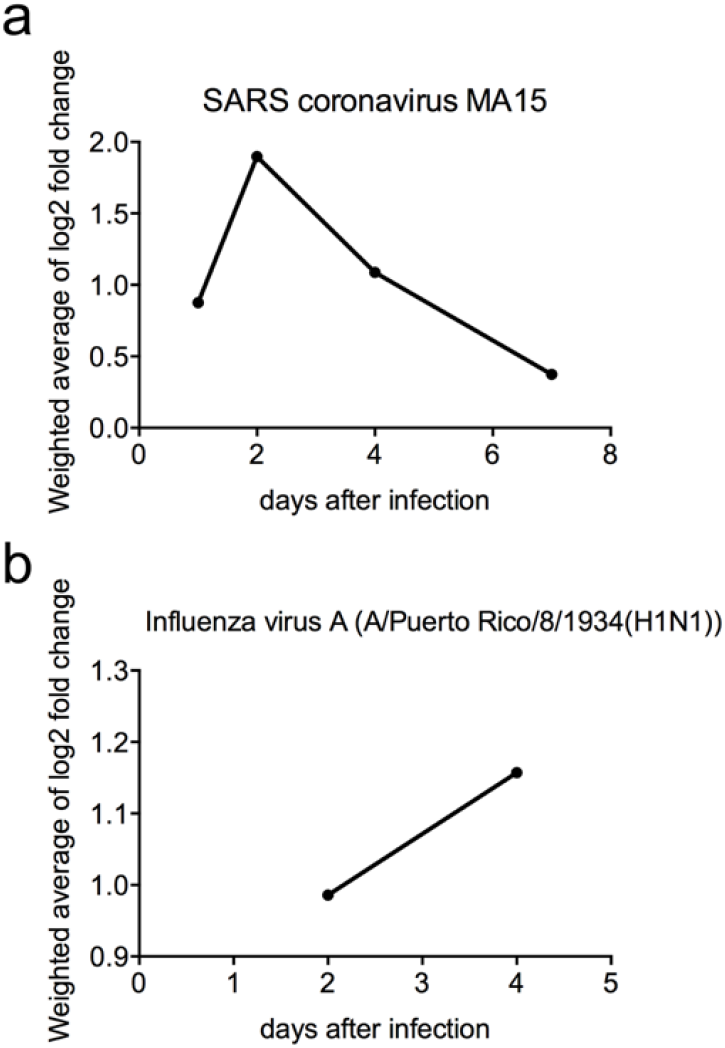
Weighted average of Tacstd2 log_2_-fold change after infection of mice with **A)** SARS coronavirus MA15 (n = 155) and **B)** Influenza A virus (n = 42) at different time points. Only data from mice infected with 10^5^ PFU in case of SARS coronavirus and 10^2^ PFU in case of Influenza A virus are shown. Data from infection with mutant viruses were excluded.

**Table 2.**
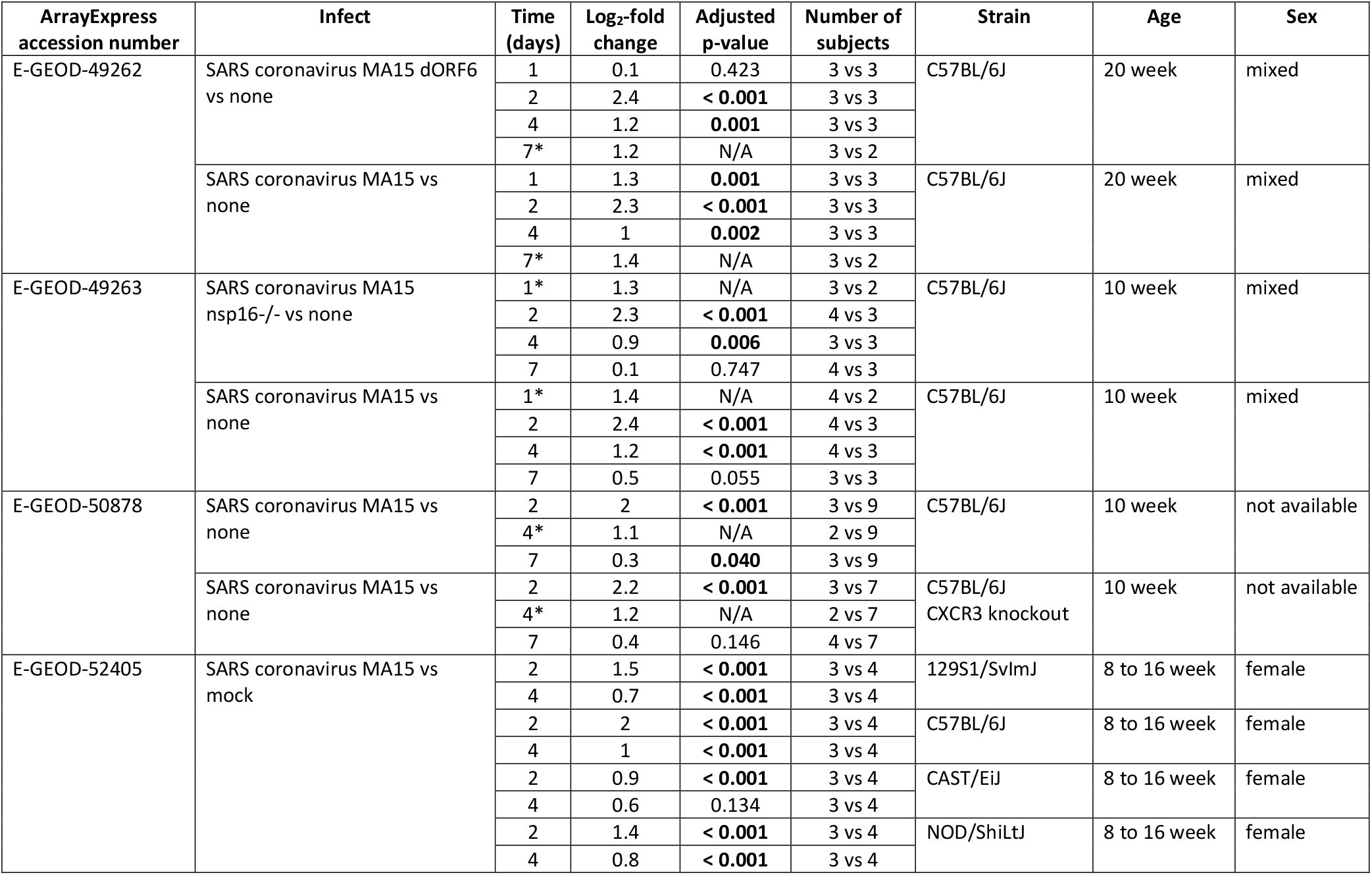
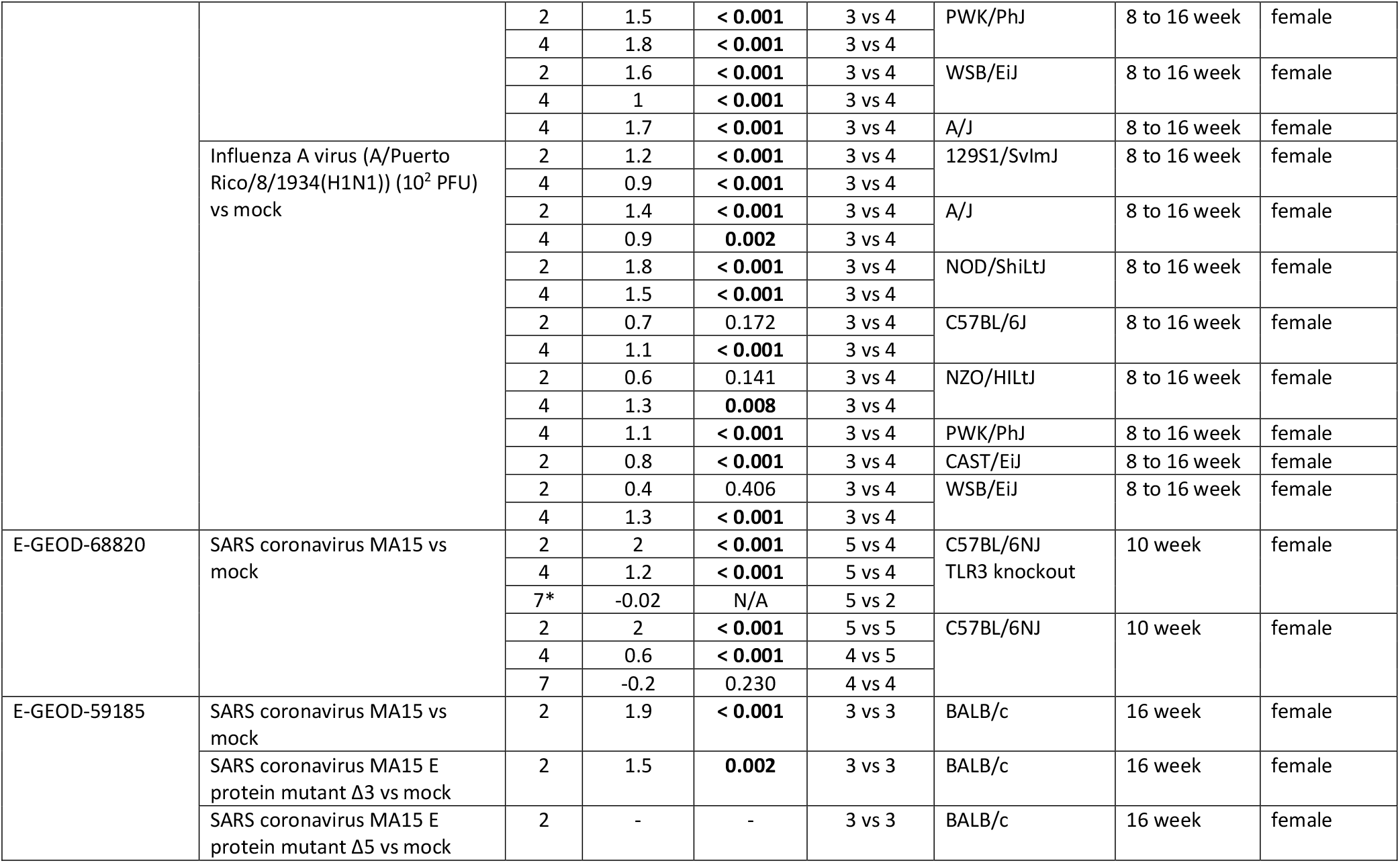
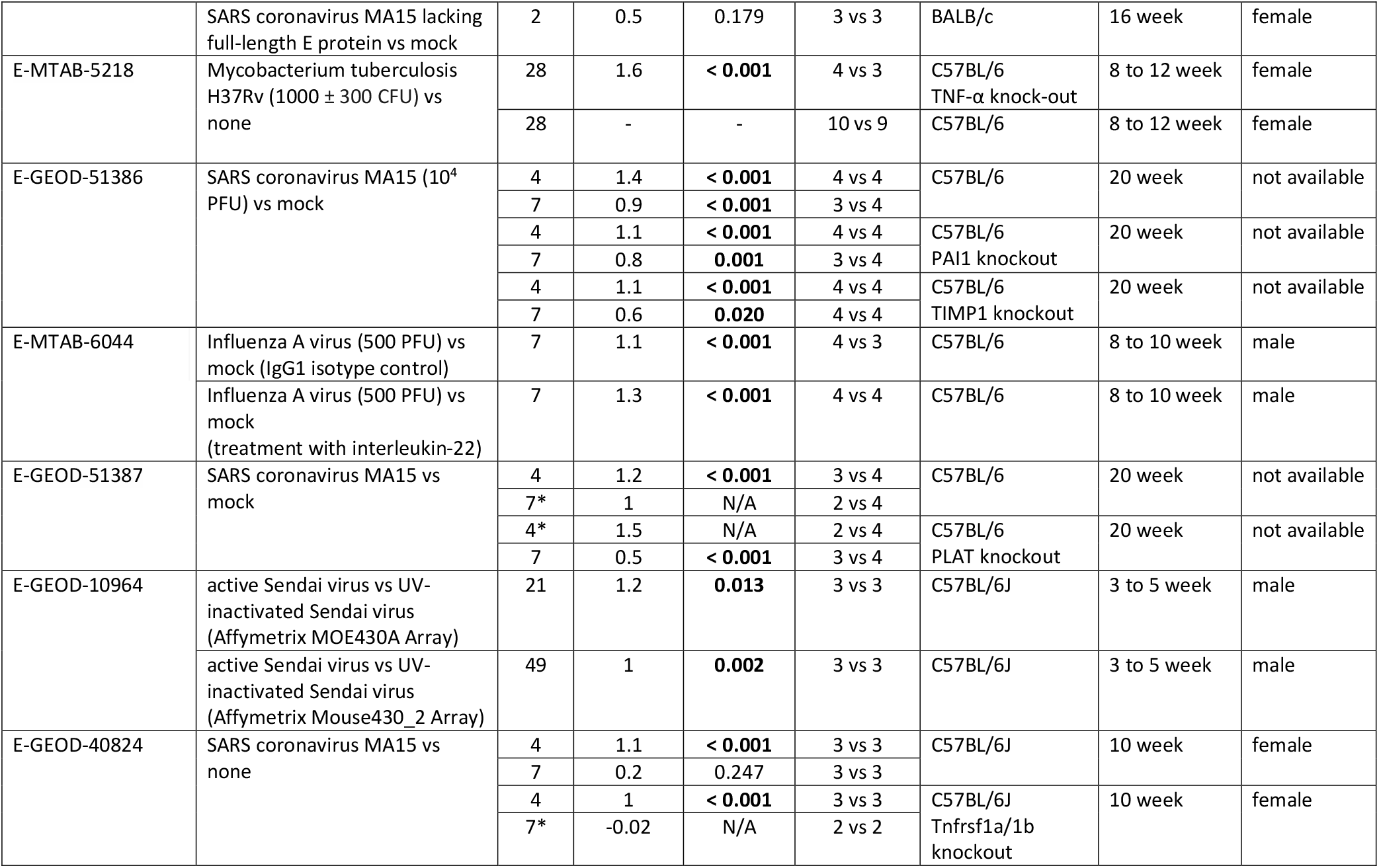
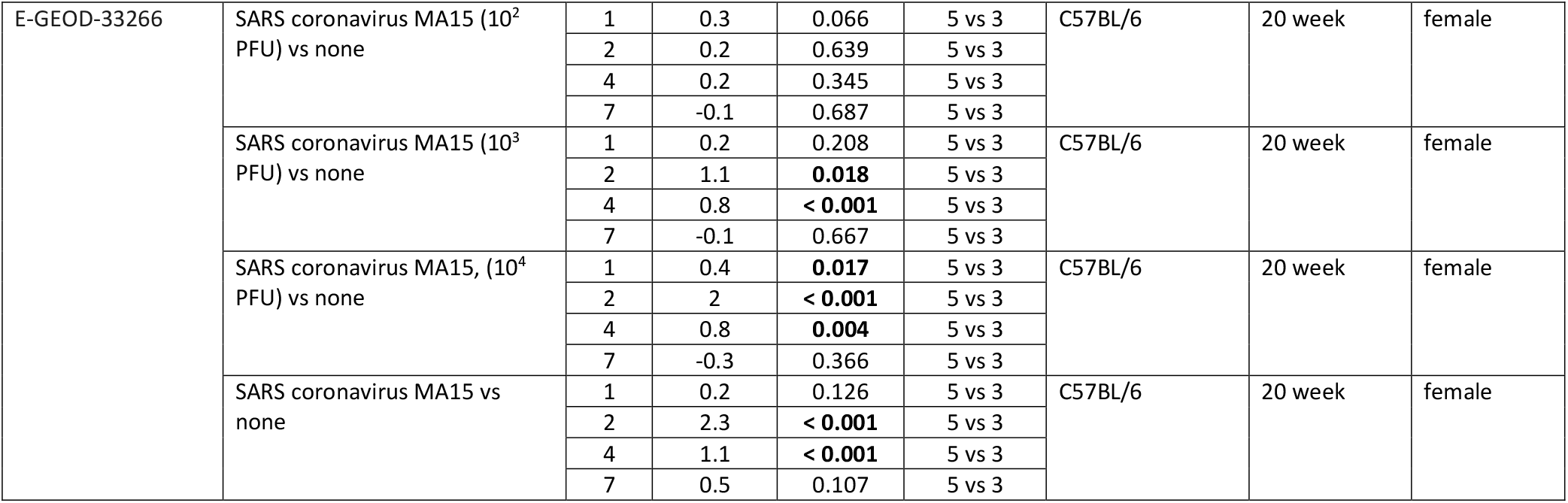
Differential Tacstd2 expression in mice after infection with various pathogens. Where not otherwise specified, viral infection dose was 10^5^ plaque forming units (PFU). Significant results (adjusted p-value < 0.05) are labeled in bold. * means that this entry was completely missing from Expression Atlas and log_2_-fold change was calculated from GEO database data using GEO2R. N/A means that p-value could not be calculated due to small number of subjects.

Interestingly, two datasets that tested *Tacstd2* expression after a longer time (**Table 2**). Lungs of mice infected with *Mycobacterium tuberculosis* H37Rv were analyzed after 28 days. In wild type C57BL/6 mice, there was no difference in *Tacstd2* expression levels in comparison to their level prior to infection, which is in agreement with other studies showing that *Tacstd2* is upregulated early after infection. However, when TNF-α knockout mice were infected, an enhanced bacterial burden, high inflammation, oedema, necrosis and increased *Tacstd2* level were detected in the lungs after 28 days. The second dataset evaluated transcriptome in mice infected with the Sendai virus. Interestingly, in this case *Tacstd2* was also upregulated late post-infection (21 and 49 days). Although, at the first sight, it might seem inconsistent with other studies, mice from this dataset developed chronic airway disease similar to human chronic airway diseases, such as asthma and COPD (40). Thus, the long upregulation of *Tacstd2* reflects the known upregulation of *TACSTD2* in human COPD (41), possibly triggered by chronic lung damage.

Significant upregulation of *TACSTD2* expression was also found in bronchoalveolar lavage cells in patients with transplanted lungs colonized by *Aspergillus fumigatus* (E-MTAB-6040). This dataset did not fulfill our inclusion criteria since it did not provide information about lung transcriptome. However, this result suggests that *TACSTD2* is upregulated after fungal infection as well (Supplementary Table S1).

The single differential expression dataset (E-GEOD-19743) containing information about *TACSTD2* levels after an injury did not fulfill inclusion criteria as it studied transcripts in blood and not lungs. However, it is interesting to note that it showed significantly upregulated *TACSTD2* in leukocytes after burn injury in both children (60 subjects) and adults (57 subjects). More details are available in Table S2 in Supplementary data. Interestingly, levels of *TACSTD2* were higher in the middle stage (11-49 days) than in early stage (<11 days) of the healing process after injury.

### TACSTD2 is upregulated in lung epithelial cells after infection

As mentioned earlier, *TACSTD2* is expressed both in lung epithelial and immune cells. It is not clear if upregulation in infected lungs is caused by increased infiltration of immune cells to the lungs or by a direct upregulation in lung epithelial cells (LECs). In order to clarify this issue, we searched GEO datasets for information about *TACSTD2* expression after infection in specific cell types.

Datasets examining expression after infecting mice with influenza virus X31 (42) (GSE148709, **Figure 5A**), and *Streptococcus pneumoniae* (43) (GSE71623, **Figure 5B**) showed that *Tacstd2* is significantly upregulated in sorted LECs when compared to uninfected cells. LECs were sorted according to their EpCAM^+^CD31^-^CD45^-^ (GSE148709) or EpCAM^+^CD45^-^ (GSE71623) expression. Upregulation of *Tacstd2* was detected also in LECs of influenza virus X31 infected *Ifnlr1^-/-^* mice.

**Figure 5.**
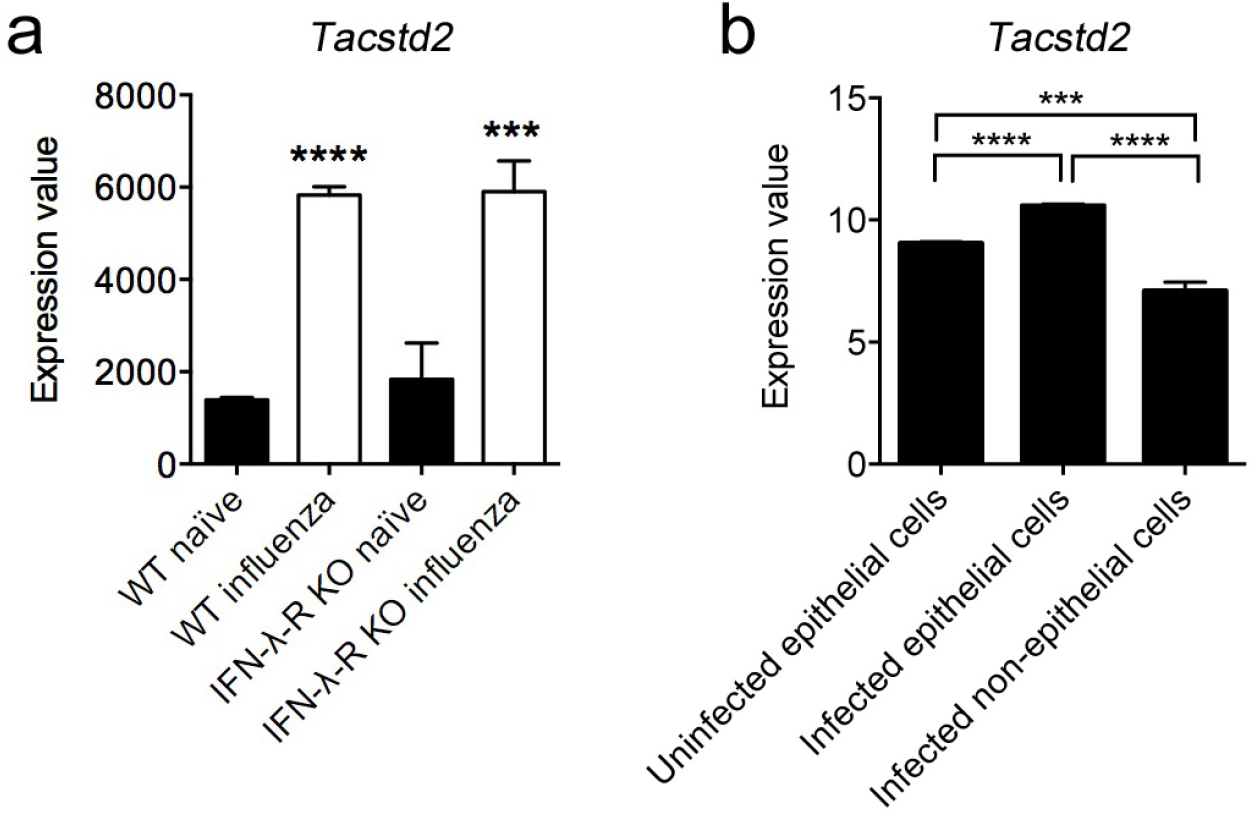
Tacstd2 expression in **A)** sorted EpCAM^+^CD31^-^CD45^-^ LECs from wt and Ifnlr^-/-^ mice infected or not with influenza virus X31 at 8 days after infection (GSE148709) (42), and **B)** sorted EpCAM^+^D45^-^ LECs from mice infected or not with Streptococcus pneumoniae at 15 hours after infection (GSE71623) (43).

Recent public dataset GSE165299 contains information about *Tacstd2* expression in mouse lung resident cells. Mice were infected with influenza viruses and 3 days post-infection alveolar epithelial cells (AECs), club cells, dendritic cells (DC), mast cells, macrophages, eosinophils, and neutrophils were isolated and the transcriptome was analyzed by high throughput sequencing. Interestingly, after infection with influenza virus, *Tacstd2* expression was increased in club cells and slightly in AECs and eosinophils, but decreased in neutrophils (**Figure 6A**). According to these data neutrophils of uninfected lungs have the highest basal expression of *Tacstd2*.

**Figure 6.**
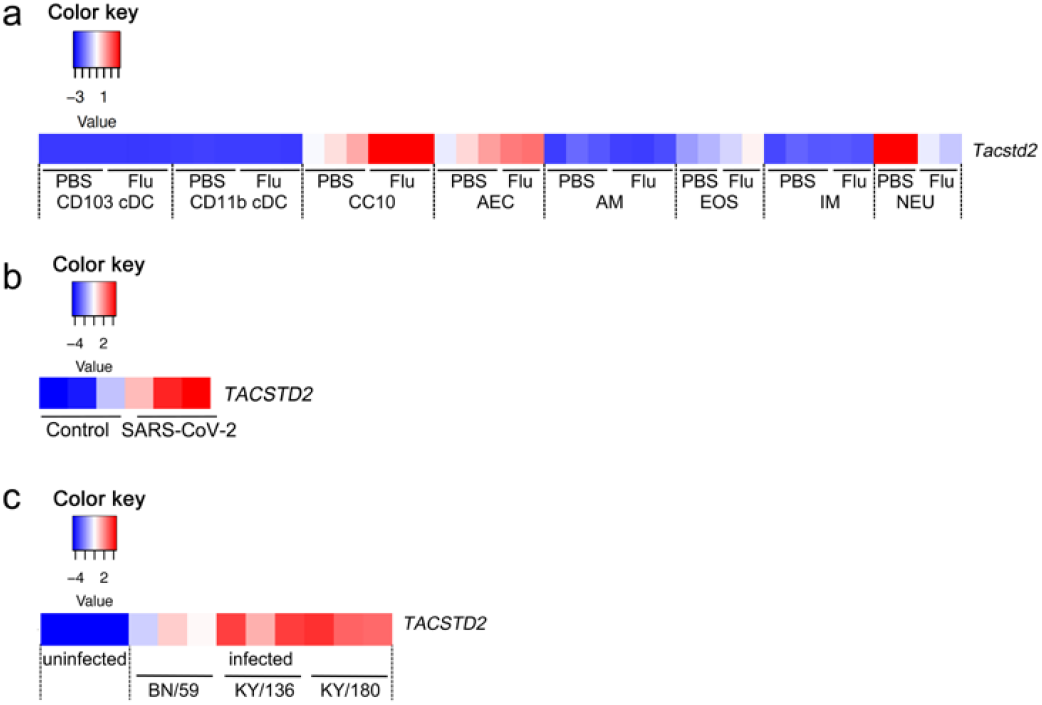
Heat map of TACSTD2/Tacstd2 expression **A)** in lung resident cells isolated from mouse lungs at 3 days post-infection (GSE165299). CD103+ cDC and CD11b+ cDC stay for dendritic cells. CC10 stays for club cells. AEC stays for alveolar epithelial cells. AM stays for alveolar macrophages. IM stays for intersticial macrophages. EOS stays for eosinophils, and NEU stays for neutrophils. **B)** in human alveolar organoids at 48 hours after infection with SARS-CoV-2 (GSE152586) (44). **C)** in human bronchial LECs at 36 hours after infection with various Influenza A H1N1 isolates (seasonal H1N1 BN/59, pandemic H1N1 KY/136 (non-fatal cases) and pandemic H1N1 KY/180 (fatal cases) (GDS4855) (45).

To further confirm the effect of infection on *TACSTD2* upregulation in lung cells we searched for datasets examining *TACSTD2* expression in LECs *in vitro*. *TACSTD2* was upregulated in human alveolar type II cell organoids infected with SARS coronavirus 2 (44) (GSE152586, **Figure 6B**) as well as in well-differentiated primary human bronchial LECs infected with various influenza A isolates (45) (GDS4855, **Figure 6C**) when compared to uninfected cells. These data confirm that LECs contribute to the increase of *TACSTD2* levels in lungs following infection.

Overall, overexpression of *TACSTD2* is an early event in the lungs challenged with various infection agents. Although increased immune cell infiltration may be partially responsible for this increase, transcriptomic studies in epithelial cells sorted from lungs of infected mice and *in vitro* infected LECs clearly proved direct upregulation of *TACSTD2*.

## Discussion

In this article, we have analysed available transcriptomic data to decipher the function of *TACSTD2* in lung tissue. Our findings may be summarized as follows. First, we have presented strong evidence that *TACSTD2* is expressed in healthy lungs of various species suggesting its evolutionarily conserved role. Second and most importantly, our analyses have shown that *Tacstd2* is significantly upregulated in mouse lungs following infection with various respiratory pathogens, suggesting its involvement in the healing process and/or immune reactions. Additionally, in most datasets, *Tacstd2* overexpression peaks early and decreases over time, suggesting it is an early reaction to infection. In case of chronic inflammation, however, *Tacstd2* remains overexpressed for long time period. The data also show that the level of *Tacstd2* upregulation depend on infection dose. Third, *Tacstd2* upregulation in the lungs after an infection is caused by a direct upregulation in LECs although some contribution of immune cells infiltrating infected lungs cannot be excluded.

Bacterial and viral pathogens are known violators of the airway epithelial barrier’s integrity, which is the first line of defense against infection, decreasing the expression, disrupting or redistributing tight and adherens junctions proteins (46–49). Such disruption significantly contributes to the pathogenesis of pulmonary infections (48). Trop2 was previously linked with maintenance of epithelial barrier function in the cornea. *TACSTD2* knockdown in corneal epithelial cells leads to decreased expression and changed subcellular localization of claudin 1, 4, 7, ZO1, and occludin which results in impaired function of corneal epithelial barrier while transduction of *TACSTD2* restored their expression and epithelial barrier function (50,51). Decreased expression and altered localization of these proteins were also observed in the corneas of GDLD patients (50). Moreover, forced expression of Trop2 can at least partially stabilize claudins and restore epithelial barrier in mouse model of congenital tufting enteropathy (52). Therefore, a possible explanation of *TACSTD2* upregulation following infection is that it represents a mechanism that helps maintain/restore the airway epithelial barrier function.

It has been shown that Trop2 is expressed in many organs during embryonal development, including lungs (9–15). It usually marks progenitor cells with high proliferation/self-renewal capacity. In fetal ovine and rat lungs, Trop2 expression positively correlates with proliferation rate (10). The cells exhibiting high Trop2 expression significantly contribute to tissue regeneration in stomach and endometrium (12,22). In stomach, transcriptome analysis further indicated that Trop2^+^ cells involved in epithelial regeneration overexpress genes that are part of a fetal developmental program (12). We therefore hypothesize that the upregulation of Trop2 in (sub)population of LECs/progenitor cells may also enhance their proliferative/pro-regenerative capacity and contribute to healing process in infected lungs. It should be noted, however, that the long term Trop2 overexpression associated with inflammation may result in hyperplasia of airway epithelium as observed in lungs of COPD patients (41).

Besides the importance of Trop2 for proper localization and function of claudins and occludins in tight junctions and possible pro-regenerative capacity of Trop2-overexpressing cells in airway epithelium, our knowledge about the role of *TACSTD2*/Trop2 in healthy tissues and during infection challenge remains limited. In cancer cells, high-throughput proteomic analysis revealed prosurvival PI3K/Akt as a major cellular signaling pathway stimulated by Trop2 (53). This has been confirmed subsequently in various cancer models (54–57), and stem cells (21,23). Activation of the Akt kinase signaling by Trop2 upregulation in response to infection may therefore enhance lung cell survival and decrease tissue damage. It should be noted, however, that controversial role of Akt kinase in modulating infection and inflammation in lungs has been reported (58,59) vs (60–63). Interestingly, both Trop2-related functional targets, tight junctions proteins and Akt kinase, were reported to be hijacked by diverse viruses to promote their infection in various tissues. While tight junction proteins may participate in regulation of viral entry, replication, dissemination and progress (64–66), activation of Akt kinase may represent a strategy of viruses to slow down apoptosis of host cells, thus prolong viral replication and enhance viral transcription (63). Thorough analysis of *Tacstd2* knockout mice upon infections with various pathogens is therefore needed to help clarify the exact role of the Trop2 protein and its signaling in lung tissue response to infections.

Besides Trop2, epithelial cell adhesion molecule (EpCAM, encoded by *TACSTD1* gene) represents another member of *TACSTD* gene family (3). Both proteins share similarities in amino acid sequence, domain structure (67), processing, cell signaling and protein interaction partners (1,68). Although both proteins interact with tight junction proteins (50,69) and participate in maintenance of epithelial barrier (51,70,71), recent study in EpCAM null mice with forced expression of Trop2 revealed that the function of both proteins is similar but not equivalent (52). Interestingly, we did not find a similar pattern of deregulation of *TACSTD1* expression in lungs after infection as observed for *TACSTD2* (Supplementary Table S3). This finding highlights an important distinction in regulation of expression of both genes and indicates another possible difference in their function.

Taken together, using available transcriptomic datasets we demonstrate that *TACSTD2* expression is evolutionarily conserved in the lungs of vertebrates and that the major source of *TACSTD2* transcript are lung epithelial cells and their progenitors. We found that lung levels of *TACSTD2* consistently increase as an early reaction to infection with various respiratory pathogens. Although this increase may represent a mechanism to maintain/restore epithelial barrier function and to mark pro-regenerative activation of progenitor cells in infected lungs, further studies are needed to clarify the exact role of the Trop2 protein and its signaling during course of lung infections and healing process.

## Acnowledgement

This work was supported by the by Ministry of Health of the Czech Republic grant NV18-07-00073, Czech Science Foundation grants no. 21-11585S, 18-001145S, Masaryk University grants MUNI/A/1689/2020, MUNI/A/1522/2020 and by the European Regional Development Fund - Project ENOCH (No. CZ.02.1.01/0.0/16_019/0000868).

