## Supplementary files for "TACSTD2 upregulation is an early reaction to lung infection"

### Supplementary Data S1: *TACSTD2* expression in lungs

#### Human (*Homo sapiens*)

#### Expression Atlas

https://www.ebi.ac.uk/gxa/about.html

| Transcriptomics | | | |
|---|---|---|---|
| **ArrayExpress Accession number** | **Developmental stage** | **Expression level (median TPM)** | **Number of subjects** |
| E-MTAB-3358 | fetal | 98 | 1 |
| | adult | 138 | 1 |
| E-MTAB-2836 | - | 228 | 5 |
| E-MTAB 1733 | - | 222 | 5 |
| E-MTAB-513 | - | 200 | 1 |
| E-MTAB-4344 | - | 187 | 1 |

| Proteomics | | |
|---|---|---|
| **PRIDE or ArrayExpress Accession number** | **Expression level** | **Number of subjects** |
| PXD010154 | 46946 | 1 |
| E-PROT-3 | below cutoff | 2 |
| E-PROT-1 | 0.00005 | 1 |

#### GTEx Portal

https://www.gtexportal.org/home/gene/TACSTD2

| **Data source** | **Expression level (median TPM)** | **Number of subjects** |
|---|---|---|
| GTEx Analysis Release V8 (dbGaP Accession phs000424.v8.p2) | 118.7 | 578 |

#### The Human Protein Atlas

https://www.proteinatlas.org/ENSG00000184292-TACSTD2/tissue/lung

| | **Expression level (median TPM)** | **Number of subjects** |
|---|---|---|
| HPA RNA Seq | 99.9 | 9 |

### Mouse (*Mus musculus*)

#### Expression Atlas

https://www.ebi.ac.uk/gxa/about.html

| Transcriptomics | | | |
|---|---|---|---|
| **ArrayExpress Accession number** | **Developmental stage or strain (where different)** | **Expression level (median TPM)** | **Number of subjects** |
| E-MTAB-3579 | embryonic day 12 | below cutoff | 1 |
| | embryonic day 14 | 2 | 1 |
| | embryonic day 15 | 2 | 1 |
| | embryonic day 16 | 3 | 1 |
| | embryonic day 17 | 8 | 1 |
| | embryonic day 18 | 9 | 1 |
| | neonate | 11 | 5 |
| | juvenile | 19 | 2 |
| | adult | 20 | 1 |
| E-MTAB-2801 | CD1 | 68 | 1 |
| | DBA/2J | 86 | 1 |
| | C57BL/6 | 93 | 1 |
| E-GEOD-74747 | - | 111 | 1 |
| E-MTAB-599 | - | 115 | 6 |
| E-MTAB-8573 | - | 112 | 3 |

| Proteomics | | |
|---|---|---|
| **DOI** | **Expression level** | **Number of subjects** |
| 10.1074/mcp.m112.024919 | below cutoff | 1 |

### Other organisms

#### Expression Atlas - transcriptomics

https://www.ebi.ac.uk/gxa/about.html

**Cattle (*Bos Taurus*)**

| ArrayExpress Accession number | Expression level (median TPM) | Number of subjects |
|---|---|---|
| E-MTAB-2798 | 119 | 3 |
| E-MTAB-2596 | 84 | 1 |

**Chicken (*Gallus gallus*)**

| ArrayExpress Accession number | Expression level (median TPM) | Number of subjects |
|---|---|---|
| E-MTAB-2797 | 2 | 3 |

**Sheep (*Ovis aries*)**

| ArrayExpress Accession number | Sex and developmental stage | Expression level (median TPM) | Number of subjects |
|---|---|---|---|
| E-MTAB-3838 | male, adult | 17 | 1 |
| | female, adult | 9 | 1 |
| | female, juvenile | 18 | 1 |
| E-GEOD-56643 | - | 66 | 1 |

**Olive baboon (*Papio anubis*)**

| ArrayExpress Accession number | Expression level (median TPM) | Number of subjects |
|---|---|---|
| E-MTAB-2848 | 38 | 1 |

**Rat (*Rattus norvegicus*)**

| ArrayExpress Accession number | Sex and developmental stage or strain (where different) | Expression level (median TPM) | Number of subjects |
|---|---|---|---|
| E-GEOD-53960 | male, juvenile | 31 | 4 |
| | male, adolescent | 42 | 4 |
| | male, adult | 49 | 4 |
| | male, elderly | 55 | 4 |
| | female, juvenile | 37 | 4 |
| | female, adolescent | 36 | 4 |
| | female, adult | 55 | 4 |
| | female, elderly | 57 | 4 |
| E-MTAB-2800 | F344/Cr1 | 121 | 1 |
| | BN/SsNHsd | 51 | 1 |
| | Sprague-Dawley | 97 | 1 |

**Pig (*Sus scrofa*)**

| ArrayExpress Accession number | Sex | Expression level (median TPM) | Number of subjects |
|---|---|---|---|
| E-MTAB-5895 | male | below cutoff | 1 |
| | female | below cutoff | 1 |

### **Bgee database**

https://bgee.org/

| **Organism** | **Expression score (median)** | **Number of subjects** |
|---|---|---|
| Human (*Homo sapies*) | 97.83 | 11 |
| Mouse (*Mus musculus*) | 72.83 | 26 |
| Chimpanzee (*Pan troglodytes*) | 68.31 | 1 |
| Macaque (*Macaca mulata*) | 92.61 | 1 |
| Rat (*Rattus norvegicus*) | 89.28 | 1 |
| Cattle (*Bos taurus*) | 78.34 | 1 |
| Pig (*Sus scrofa*) | 64.33 | 2 |
| Rabbit (*Oryctolagus cuniculus*) | 96.55 | 1 |
| Opossum (*Monodelphis domestica*) | 90.48 | 1 |
| Chicken (*Gallus gallus*) | 27.63 | 1 |

**Supplementary Table S1. Differential *TACSTD2* expression of bronchoalveolar lavage cells in patients with transplanted lungs colonized by *Aspergillus fumigatus* (E-MTAB-6040)**

| Infect | $Log_2$-fold change | Adjusted p-value | Number of subjects |
|---|---|---|---|
| *Aspergillus fumigatus* vs none | 1.8 | **0.028** | 6 vs 5 |

**Supplementary Table S2. Differential *TACSTD2* expression of blood samples from pediatric and adults patients with burn injury (E-GEOD-19743**). Samples were collected in early stage (<11 days) and middle stage (11-49 days) after injury.

| | $Log_2$-fold change | Adjusted p-value | Number of subjects |
|---|---|---|---|
| Adult, early stage vs adult, control | 1 | **0.035** | 29 vs 28 |
| Adult, middle stage vs adult, control | 3 | **< 0.001** | 28 vs 28 |
| Child, early stage vs child, control | 0.7 | **0.038** | 25 vs 35 |
| Child, middle stage vs child, control | 2.6 | **< 0.001** | 24 vs 35 |

**Supplementary Table S3. Differential *Epcam* expression in mice after infection with various pathogens**. Where not otherwise specified, viral infection dose was $10^5$ plaque forming units (PFU). Significant results (adjusted p-value < 0.05) are labeled in bold. * means that this entry was missing from Expression Atlas and $\log_2$-fold change was calculated from GEO database data using GEO2R. N/A means that p-value could not be calculated due to small number of subjects or that change in *Epcam* expression was not analyzed in given dataset.

| ArrayExpress accession number | Infect | Time (days) | $\text{Log}_2$-fold change | Adjusted p-value | Number of subjects | Strain | Age | Sex |
|---|---|---|---|---|---|---|---|---|
| E-GEOD-49262 | SARS coronavirus MA15 dORF6 vs none | 1 | -0.3 | **0.034** | 3 vs 3 | C57BL/6J | 20 week | mixed |
| | | 2 | 0.2 | 0.145 | 3 vs 3 | | | |
| | | 4 | -0.1 | 0.783 | 3 vs 3 | | | |
| | | 7* | -0.6 | N/A | 3 vs 2 | | | |
| | SARS coronavirus MA15 vs none | 1 | -0.3 | 0.118 | 3 vs 3 | C57BL/6J | 20 week | mixed |
| | | 2 | - | - | 3 vs 3 | | | |
| | | 4 | 0.3 | 0.352 | 3 vs 3 | | | |
| | | 7* | -0.5 | N/A | 3 vs 2 | | | |
| E-GEOD-49263 | SARS coronavirus MA15 nsp16-/- vs none | 1* | -0.4 | N/A | 3 vs 2 | C57BL/6J | 10 week | mixed |
| | | 2 | 0.4 | **0.005** | 4 vs 3 | | | |
| | | 4 | -0.3 | 0.132 | 3 vs 3 | | | |
| | | 7 | -0.6 | **0.007** | 4 vs 3 | | | |
| | SARS coronavirus MA15 vs none | 1* | -0.4 | N/A | 4 vs 2 | C57BL/6J | 10 week | mixed |
| | | 2 | 0.2 | 0.131 | 4 vs 3 | | | |
| | | 4 | -0.1 | 0.674 | 4 vs 3 | | | |
| | | 7 | -0.8 | **< 0.001** | 3 vs 3 | | | |
| E-GEOD-50878 | SARS coronavirus MA15 vs none | 2 | 0.3 | 0.056 | 3 vs 9 | C57BL/6J | 10 week | not available |
| | | 4* | -0.3 | N/A | 2 vs 9 | | | |
| | | 7 | -0.4 | **0.007** | 3 vs 9 | | | |
| | SARS coronavirus MA15 vs none | 2 | 0.2 | **0.016** | 3 vs 7 | C57BL/6J CXCR3 knockout | 10 week | not available |
| | | 4* | 0.4 | N/A | 2 vs 7 | | | |
| | | 7 | -0.7 | **< 0.001** | 4 vs 7 | | | |
| E-GEOD-52405 | SARS coronavirus MA15 vs mock | 2 | 0.4 | 0.097 | 3 vs 4 | 129S1/SvImJ | 8 to 16 week | female |
| | | 4 | - | - | 3 vs 4 | | | |

| | | | | | | | | |
|---|---|---|---|---|---|---|---|---|
| | | 2 | 0.3 | 0.094 | 3 vs 4 | C57BL/6J | 8 to 16 week | female |
| | | 4 | 0.3 | 0.067 | 3 vs 4 | | | |
| | | 2 | 0.4 | **0.039** | 3 vs 4 | CAST/EiJ | 8 to 16 week | female |
| | | 4 | 0.5 | **0.043** | 3 vs 4 | | | |
| | | 2 | 0.2 | 0.574 | 3 vs 4 | NOD/ShiLtJ | 8 to 16 week | female |
| | | 4 | 0.1 | 0.798 | 3 vs 4 | | | |
| | | 2 | 0.5 | **0.011** | 3 vs 4 | PWK/PhJ | 8 to 16 week | female |
| | | 4 | 0.6 | **< 0.001** | 3 vs 4 | | | |
| | | 2 | 0.3 | 0.297 | 3 vs 4 | WSB/EiJ | 8 to 16 week | female |
| | | 4 | 0.5 | **0.013** | 3 vs 4 | | | |
| | | 4 | 0.3 | 0.191 | 3 vs 4 | A/J | 8 to 16 week | female |
| | Influenza A virus (A/Puerto Rico/8/1934(H1N1)) ($10^2$ PFU) vs mock | 2 | 0.4 | 0.085 | 3 vs 4 | 129S1/SvImJ | 8 to 16 week | female |
| | | 4 | 0.3 | 0.149 | 3 vs 4 | | | |
| | | 2 | 0.2 | 0.474 | 3 vs 4 | A/J | 8 to 16 week | female |
| | | 4 | 0.2 | 0.802 | 3 vs 4 | | | |
| | | 2 | 0.9 | **< 0.001** | 3 vs 4 | NOD/ShiLtJ | 8 to 16 week | female |
| | | 4 | 0.3 | 0.068 | 3 vs 4 | | | |
| | | 2 | -0.1 | 0.93 | 3 vs 4 | C57BL/6J | 8 to 16 week | female |
| | | 4 | 0.5 | **0.025** | 3 vs 4 | | | |
| | | 2 | 0.2 | 0.626 | 3 vs 4 | NZO/HlLtJ | 8 to 16 week | female |
| | | 4 | 0.5 | 0.24 | 3 vs 4 | | | |
| | | 4 | - | - | 3 vs 4 | PWK/PhJ | 8 to 16 week | female |
| | | 2 | 0.3 | 0.232 | 3 vs 4 | CAST/EiJ | 8 to 16 week | female |
| | | 2 | - | - | 3 vs 4 | WSB/EiJ | 8 to 16 week | female |
| | | 4 | 0.7 | **< 0.001** | 3 vs 4 | | | |
| E-GEOD-68820 | SARS coronavirus MA15 vs mock | 2 | 0.3 | **0.006** | 5 vs 4 | C57BL/6NJ TLR3 knockout | 10 week | female |
| | | 4 | 0.2 | **0.022** | 5 vs 4 | | | |
| | | 7* | -0.02 | N/A | 5 vs 2 | | | |
| | | 2 | 0.2 | **0.034** | 5 vs 5 | C57BL/6NJ | 10 week | female |
| | | 4 | 0.2 | **0.015** | 4 vs 5 | | | |
| | | 7 | -0.2 | **0.014** | 4 vs 4 | | | |

| | | | | | | | | |
|---|---|---|---|---|---|---|---|---|
| E-GEOD-59185 | SARS coronavirus MA15 vs mock | 2 | N/A | N/A | 3 vs 3 | BALB/c | 16 week | female |
| | SARS coronavirus MA15 E protein mutant Δ3 vs mock | 2 | N/A | N/A | 3 vs 3 | BALB/c | 16 week | female |
| | SARS coronavirus MA15 E protein mutant Δ5 vs mock | 2 | N/A | N/A | 3 vs 3 | BALB/c | 16 week | female |
| | SARS coronavirus MA15 lacking full-length E protein vs mock | 2 | N/A | N/A | 3 vs 3 | BALB/c | 16 week | female |
| E-MTAB-5218 | Mycobacterium tuberculosis H37Rv (1000 ± 300 CFU) vs none | 28 | - | - | 4 vs 3 | C57BL/6 TNF-α knock-out | 8 to 12 week | female |
| | | 28 | -0.2 | **< 0.001** | 10 vs 9 | C57BL/6 | 8 to 12 week | female |
| E-GEOD-51386 | SARS coronavirus MA15 ($10^4$ PFU) vs mock | 4 | 0.1 | 0.347 | 4 vs 4 | C57BL/6 | 20 week | not available |
| | | 7 | -0.2 | **0.006** | 3 vs 4 | | | |
| | | 4 | 0.2 | 0.185 | 4 vs 4 | C57BL/6 PAI1 knockout | 20 week | not available |
| | | 7 | -0.3 | **0.026** | 3 vs 4 | | | |
| | | 4 | -0.1 | 0.405 | 4 vs 4 | C57BL/6 TIMP1 knockout | 20 week | not available |
| | | 7 | -0.5 | **< 0.001** | 4 vs 4 | | | |
| E-MTAB-6044 | Influenza A virus (500 PFU) vs mock (treatment with IgG1 isotype control) | 7 | N/A | N/A | 4 vs 3 | C57BL/6 | 8 to 10 week | male |
| | Influenza A virus (500 PFU) vs mock (treatment with interleukin-22) | 7 | N/A | N/A | 4 vs 4 | C57BL/6 | 8 to 10 week | male |
| E-GEOD-51387 | SARS coronavirus MA15 vs mock | 4 | 0.2 | **0.033** | 3 vs 4 | C57BL/6 | 20 week | not available |
| | | 7* | -0.4 | N/A | 2 vs 4 | | | |
| | | 4* | - | - | 2 vs 4 | C57BL/6 PLAT knockout | 20 week | not available |
| | | 7 | -0.3 | **0.006** | 3 vs 4 | | | |
| E-GEOD-10964 | active Sendai virus vs UV-inactivated Sendai virus (Affymetrix MOE430A Array) | 21 | 0.7 | **0.005** | 3 vs 3 | C57BL/6J | 3 to 5 week | male |

| | | | | | | | | |
|---|---|---|---|---|---|---|---|---|
| | active Sendai virus vs UV-inactivated Sendai virus (Affymetrix Mouse430_2 Array) | 49 | 0.7 | **0.010** | 3 vs 3 | C57BL/6J | 3 to 5 week | male |
| E-GEOD-40824 | SARS coronavirus MA15 vs none | 4 | -0.1 | 0.675 | 3 vs 3 | C57BL/6J | 10 week | female |
| | | 7 | -0.2 | 0.124 | 3 vs 3 | | | |
| | | 4 | - | - | 3 vs 3 | C57BL/6J Tnfrsf1a/1b knockout | 10 week | female |
| | | 7* | -0.5 | N/A | 2 vs 2 | | | |
| E-GEOD-33266 | SARS coronavirus MA15 ($10^2$ PFU) vs none | 1 | 0.3 | **0.040** | 5 vs 3 | C57BL/6 | 20 week | female |
| | | 2 | 0.5 | 0.237 | 5 vs 3 | | | |
| | | 4 | 0.1 | 0.382 | 5 vs 3 | | | |
| | | 7 | - | - | 5 vs 3 | | | |
| | SARS coronavirus MA15 ($10^3$ PFU) vs none | 1 | 0.4 | **0.009** | 5 vs 3 | C57BL/6 | 20 week | female |
| | | 2 | 0.2 | 0.444 | 5 vs 3 | | | |
| | | 4 | 0.1 | 0.506 | 5 vs 3 | | | |
| | | 7 | -0.3 | **0.020** | 5 vs 3 | | | |
| | SARS coronavirus MA15 ($10^4$ PFU) vs none | 1 | 0.6 | **0.006** | 5 vs 3 | C57BL/6 | 20 week | female |
| | | 2 | 0.2 | 0.408 | 5 vs 3 | | | |
| | | 4 | 0.3 | 0.080 | 5 vs 3 | | | |
| | | 7 | -0.3 | **0.013** | 5 vs 3 | | | |
| | SARS coronavirus MA15 vs none | 1 | 0.6 | **0.020** | 5 vs 3 | C57BL/6 | 20 week | female |
| | | 2 | 0.3 | 0.060 | 5 vs 3 | | | |
| | | 4 | 0.1 | 0.455 | 5 vs 3 | | | |
| | | 7 | -0.3 | **0.017** | 5 vs 3 | | | |

**Supplementary Fig. S1 Immunohistochemical detection of Trop2 in paraffin sections of human, mouse, and pig lung tissue.** Human lungs – positive staining in epithelium of airway and alveoli. Mouse/Pig lungs – positive staining only in basolateral parts of airway epithelium.

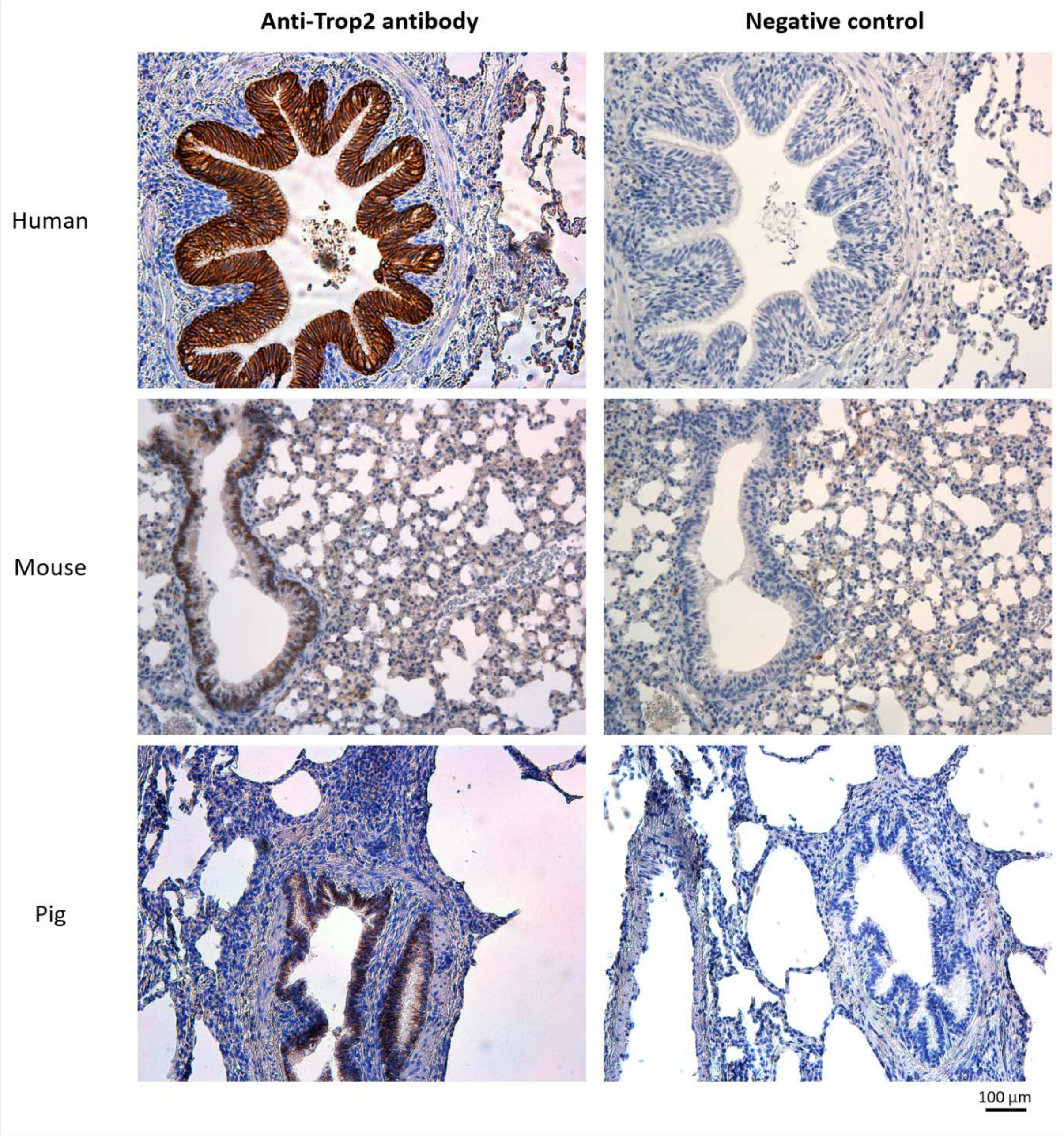


.
